# CREPE: A Shiny App for Transcription Factor Cataloguing

**DOI:** 10.1101/2022.11.14.516198

**Authors:** Diego A. Rosado-Tristani, José A. Rodríguez-Martínez

## Abstract

Transcription factors (TFs) are proteins that directly interpret the genome to regulate gene expression and determine cellular phenotypes. TF identification is a common first step in unravelling gene regulatory networks. We present CREPE, an R Shiny app to catalogue and annotate TFs. CREPE was benchmarked against curated human transcription factor datasets. Next, we use CREPE to explore the TF repertoires of *Heliconius erato and Heliconius melpomene* butterflies. CREPE is available as a Shiny app package available at GitHub (github.com/dirostri/CREPE).

## Introduction

Transcription factors (TFs) are sequence-specific DNA-binding proteins that regulate transcription. TF proteins bind to cis-regulatory elements (CREs), genomic regions that regulate the expression of nearby genes. Examples of CREs include promoters, enhancers, and silencers (Wittkopp and Kalay, 2011). By regulating spatiotemporal gene expression, TFs determine cell fate during development and cellular responses to the environment. TFs are classified into structural families based on previously described DNA-binding domains (DBD) (Weirauch and Hughes, 2011). In eukaryotes, over 80 families of TFs are recognized, and the list includes homeodomains (e.g., Hox genes), nuclear receptors (ESR1; estrogen receptor) and P53 (TP53; tumor suppressor). Additional families of TFs are believed to exist (Weirauch and Hughes, 2011). The structure of a TF contains at least one DBD. However, they can have more than one DBD and additional regulatory domains (Frietze and Farnham, 2011). Therefore, TFs and their regulation have a global effect on cellular phenotypes, highlighting the need for their identification. Advances in DNA sequencing technologies have greatly increased the number of available genome sequences, enabling researchers to interrogate genetic and molecular mechanisms in their favorite systems (Hotaling et al., 2021). A typical goal is to unravel gene regulatory networks controlling development in a biological system, and a common first step is to identify which TFs are expressed in specific tissues. A comprehensive catalog of all TFs encoded by a genome is oftentimes missing, especially in non-model organisms. With an ever-increasing number of available genomes, there is a need for tools that can aid downstream characterization and make the most of these genomes. To address this need, we present the Cis-regulatory Element-binding Protein Elucidator (CREPE), a tool to catalog and annotate TF proteins.

## Description

CREPE performs two main functions: transcription factor 1) cataloguing and 2) annotation.

### Transcription Factor Cataloguing

TFs are classified into structural families according to their DBDs. By parsing genome assemblies for previously described DBDs, it is possible to identify putative TFs. To run the TF Cataloguing function requires a species name and a protein sequence file (.fasta). A protein domain search analysis is performed using HMMER Suite (Eddy, 2011) to scan the protein sequences against 82 eukaryotic TF DBD models (Weirauch and Hughes, 2011) (.hmm) from PFAM (Mistry et al., 2021). Next, CREPE displays a TF family distribution plot and provides a sequence file with the identified putative TFs (CREPE_FL.fasta). This file can now be used for phylogenetic inferences using the researcher’s software of choice.

### Transcription Factor Annotation

To run the TF Annotation function requires gene trees (in Newick format) of the putative TFs previously run by the user, and a metadata tab-separated file (.tsv) which cross-references a sequence id to its gene symbol. During execution, each tree is parsed to assign the putative TF the gene symbol of its closest relative in the gene tree as measured by the patristic distance using the ape 5.0 package (Paradis and Schliep, 2019). After execution, CREPE displays a table showing the resulting mappings of the putative TFs and makes them available for download.

## Benchmark

We benchmarked CREPE’s performance to manually curated human TF datasets. We applied the TF Cataloguing function of CREPE to the human proteome (GRCh38.p13, Ensembl 107) (Cunningham et al., 2022). The proteome was preprocessed to only include the primary isoform per gene [from 120,712 to 23,486 sequences], and sequences derived from alternative mappings were removed [from 23,486 to 20,409 sequences]. We identified 1519 genes (~7.4% of the input sequences) as putative TFs across 51 families (Supplementary Table 1). As expected, the five largest TF families were the C2H2 zinc fingers (zf-C2H2), homeodomains, basic helix-loop-helix (bHLH), basic leucine zippers (bZIP) and forkhead (Fig.1).

**Fig. 1.**
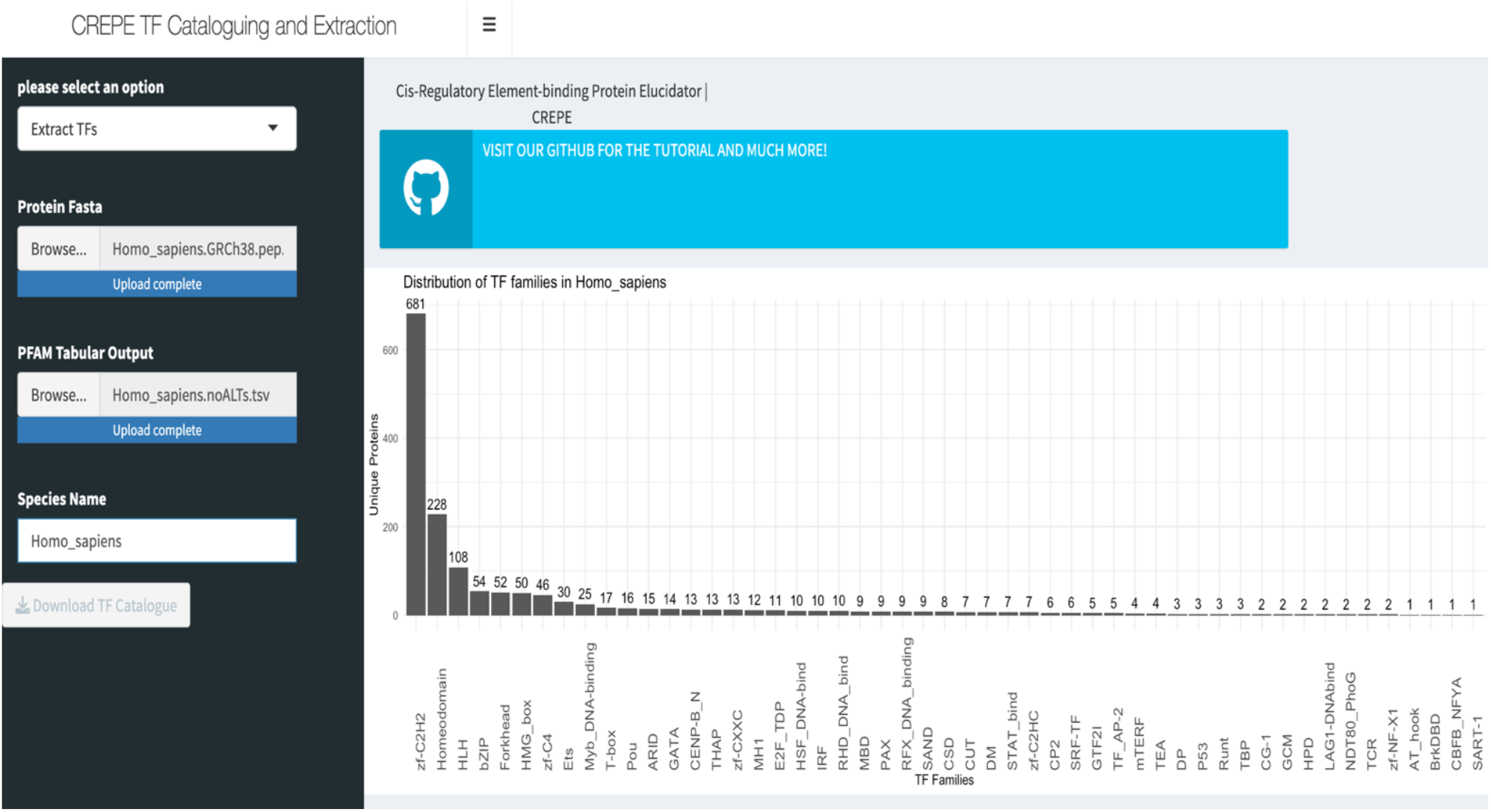
CREPE output after running the TF Catalogue function. The main panel shows the distribution of TFs by family. In this example we are using a non-redundant human proteome of 1519 human TFs.

To validate our findings, we compared them to curated human TF datasets: CisBP 2.00 (Weirauch et al., 2014), which was preprocessed to include TFs with known TF DBD [from 1639 to 1546 genes] and the Human Transcription Factors catalog (hTFcat) (Lambert et al., 2018), which was also preprocessed to remove TFs with unknown DBDs [from 1639 to 1570 genes]. CREPE retrieved 91.6% (1416/1546) of the TFs in CisBP and 90.2% (1416/1570) of the TFs in the hTFcat (Fig. 2A). Comparison by family shows that 28 of the 51 (55%) TF families identified by CREPE showed parity (Fig. 2B; Supplementary Table 2).

**Fig. 2.**
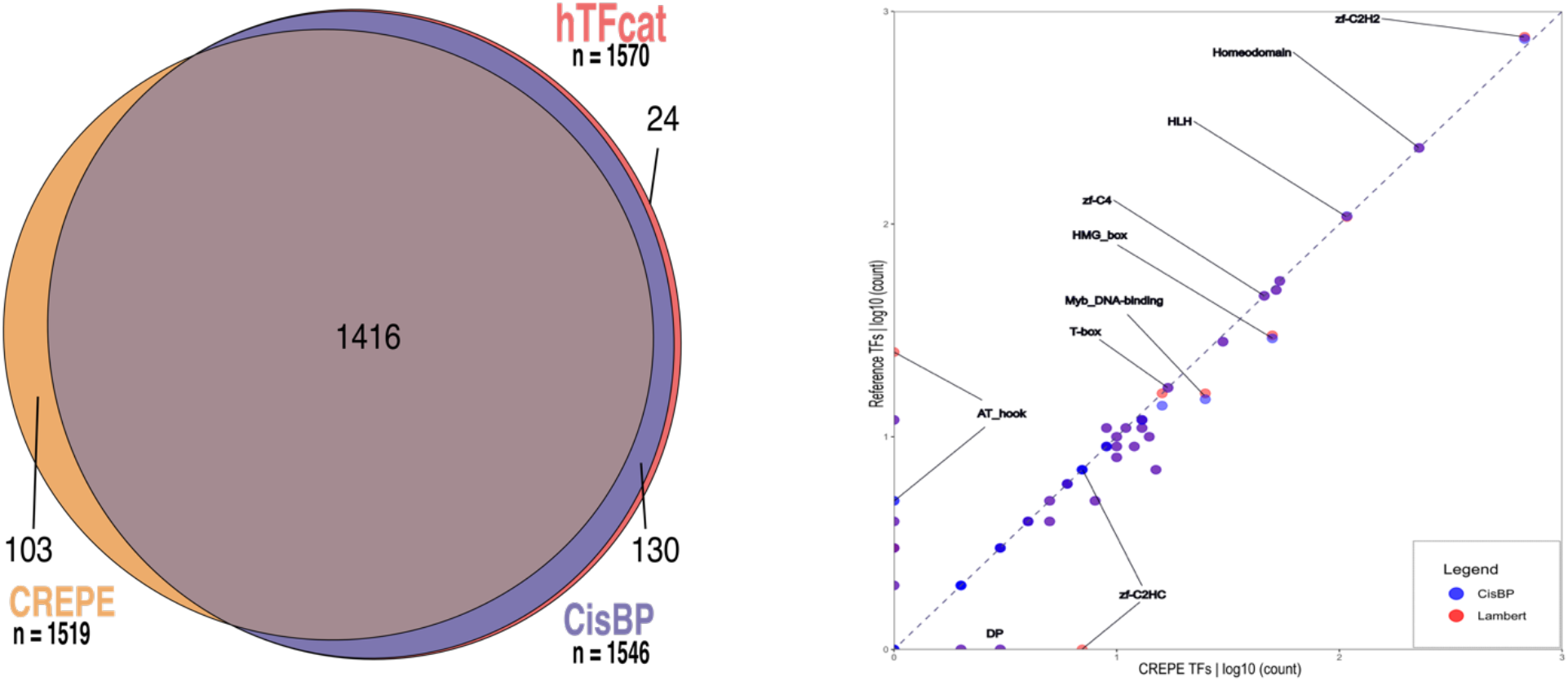
CREPE Benchmarking. A) Venn diagram showing overlap between the CREPE TF catalog (orange) against CisBP (purple) and hTFcat (red). B) Scatter plot of TFs identified by CREPE with TFs from CisBP and hTFcat by TF family; dashed line shows 1:1 relationship.

CREPE identified 103 genes as TFs across 21 families, which were not included in the reference TF datasets (Supplementary Fig. 1A; Supplementary Table 2). The largest family included in this group (21/103; 20%) are the high-mobility group box, and includes HMGB1-4, TOX, and TOX2-4 which to date, have not yielded DNA-binding specificity motifs (Lambert et al., 2018). The second largest family is the C2H2 zinc fingers with 20 genes, of which 11 are annotated by Ensembl as novel and 3 as read-throughs (Cunningham et al., 2022) (Supplementary Fig. 1B; Supplementary Table 2). CREPE also identified genes such as HSFX3, HSFX4, DUXB, FOXL3, KLF18 that are members of known TF families. Whether they possess any DNA-binding sequence-specificity and TF activity needs to be validated experimentally.

CREPE missed 130 genes that are in both reference human TF datasets, spanning 16 families (Supplementary Figure 2). The largest family in this group accounting for over half of the genes (79/130; 60.8%) are C2H2 zinc fingers. A notable difference between CREPE and the references appeared in the AT Hook and C2H2 zinc finger families. This was expected because it is known that current models can make the classification of C2H2 zinc fingers and AT Hook DBDs difficult (Lambert et al., 2018). Taken together, our strategy using CREPE to identify the human TFs is summarized with a true positive rate of 90.2%.

## Application

We applied CREPE to catalog the TF repertoire of *Heliconius melpomene* and *Heliconius erato*,two species of butterflies that are found throughout Central and South America and are a popular model system to study phenotypic determination through their wing coloration patterning mimicry (McMillan et al., 2020). We obtained predicted proteomes for both species from LepBase (Challi et al., 2016). Next, we applied the CREPE TF Cataloguing function and catalogued 664 and 599 TFs in *H. melpomene* and *H. erato*, respectively (Supplementary Figure 3). In both species, TFs were distributed across 51 families. The top three families in both butterflies are zf-C2H2, homeodomain, and bHLH, accounting for over half of the identified putative TFs. In preparation to run the CREPE TF Annotation function, we generated the required putative TF gene trees using OrthoFinder (Emms and Kelly, 2019) and 20 animal proteomes from Ensemble (Cunningham et al., 2022) (Supplementary Table 3). These gene trees were then used as inputs for the TF Annotation function. The annotations of the putative TFs for *H. erato* and *H. melpomene* mapped to *Drosophila melanogaster* are available in our GitHub.

## Conclusion

In this work, we described CREPE, a Shiny app to systematically catalogue and annotate TFs. CREPE was benchmarked against curated human TFs datasets, obtaining a 90.2% true positive rate. Using CREPE, we identified putative TFs that are currently not listed in curated references, suggesting that by using CREPE a user can obtain an updated TF catalog. The intention behind this tool is to allow researchers to explore the TFs in poorly annotated genomes or from understudied organisms. To showcase this functionality, we catalogued the TF repertoire of *H. melpomene* and *H. erato* butterflies. Taken together, CREPE provides a path forward for large-scale TF identification. A tutorial on how to execute CREPE can be found in our GitHub (github.com/dirostri/CREPE).

## Supporting information

SupportingInformation

## Acknowledgements

We would like to thank Dr. Steven van Belleghem for his insights and support in this project. We would like to thank Dr. Humberto Ortiz-Zuazaga for their help in setting up the CREPE binder launcher.

## References

Challi, R. et al. (2016) Lepbase: the Lepidopteran genome database. bioRxiv, 56994.

Cunningham, F. et al. (2022) Ensembl 2022. Nucleic Acids Research, 50, D988–95.

Eddy, S (2011) Accelerated Profile HMM Searches. PLOS Computational Biology, 7(10), e1002195.

Emms DM, Kelly S. (2019) OrthoFinder: phylogenetic orthology inference for comparative genomics. Genome Biology. 20:238.

Frietze S, Farnham P. (2011) ‘Transcription Factor Effector Domains’, in Hughes, T.R. (ed.) A Handbook of Transcription Factors. Dordrecht, Heidelberg, London, New York: Springer, 261–278.

Hotaling, S. et al. (2021) Toward a genome sequence for every animal: Where are we now?. PNAS, 118, e2109019118.

Lambert, S. et al. (2018) The Human Transcription Factors. Cell, 172, 650–665.

McMillan, W et al. (2020) From Patterning Genes to Process: Unraveling the Gene Regulatory Networks That Pattern Heliconius Wings. Frontiers in Ecology and Evolution, Vol 8, fevo.2020.00221

Mistry, J. et al. (2020) Pfam: The protein families database in 2021. Nucleic Acids Research, Vol. 49 Issue D1, D412–D419.

Paradis E, Schliep K (2019) ape 5.0: an environment for modern phylogenetics and evolutionary analyses in R. Bioinformatics, 35, 526–528.

Weirauch, M. et al. 2014) Determination and Inference of Eukaryotic Transcription Factor Sequence Specificity. Cell, 158, 1431–1443.

Weirauch M, Hughes T.R. (2011) ‘A Catalogue of Eukaryotic Transcription Factor Types, Their Evolutionary Origin, and Species Distribution’, in Hughes, T.R. (ed.) A Handbook of Transcription Factors. Dordrecht, Heidelberg, London, New York: Springer, 25–74.

Wittkopp PJ, Kalay G. (2012) Cis-regulatory elements: molecular mechanisms and evolutionary processes underlying divergence. Nature Reviews Genetics, 13:59–69.

