## SupportingInformation for "CREPE: A Shiny App for Transcription Factor Cataloguing"

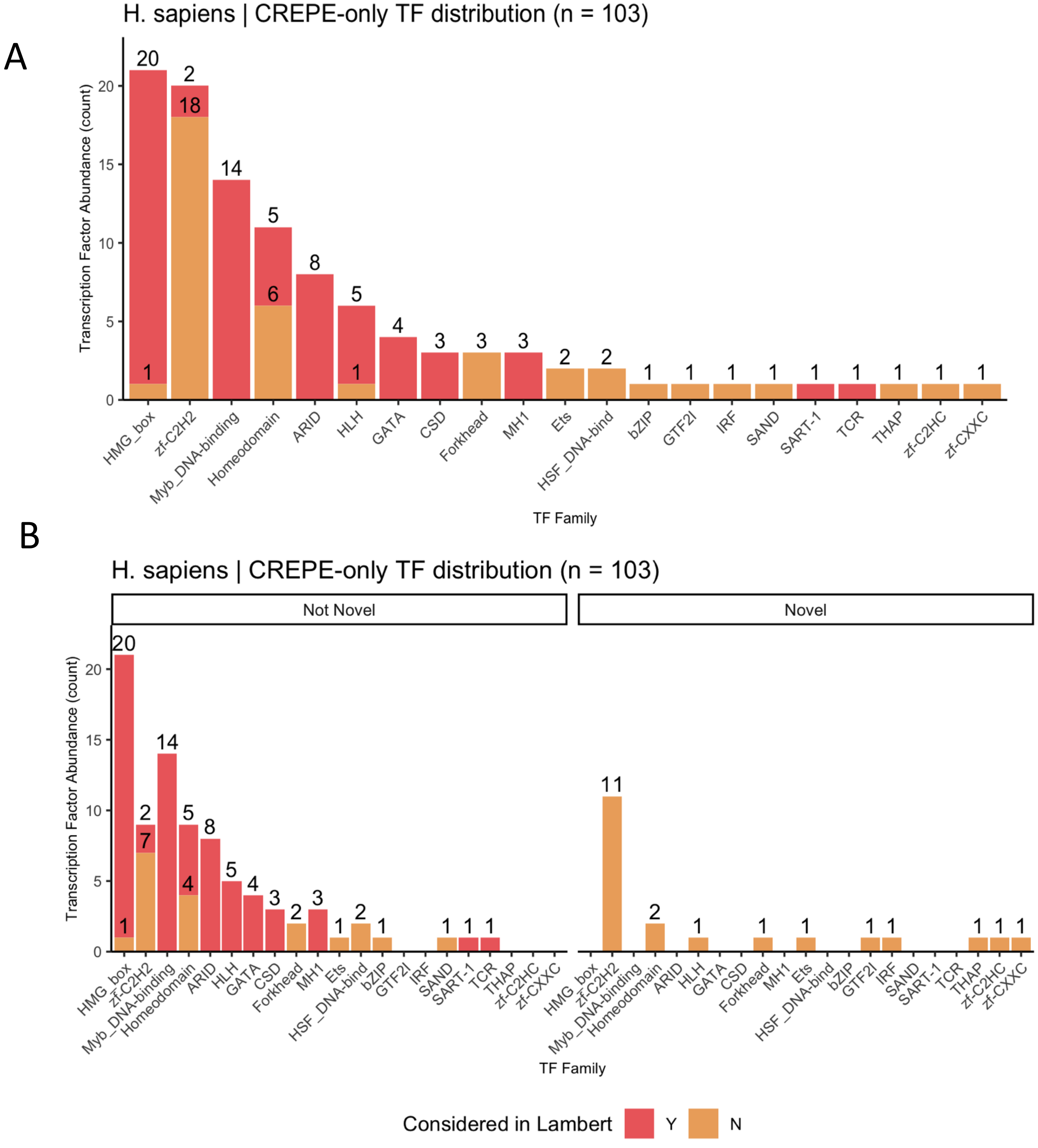


**Supplementary Figure 1:** A) Transcription factor family distribution plot of the CREPE-only group; B) subdivided into novel genes as annotated by Ensembl (Cunningham et al., 2022)

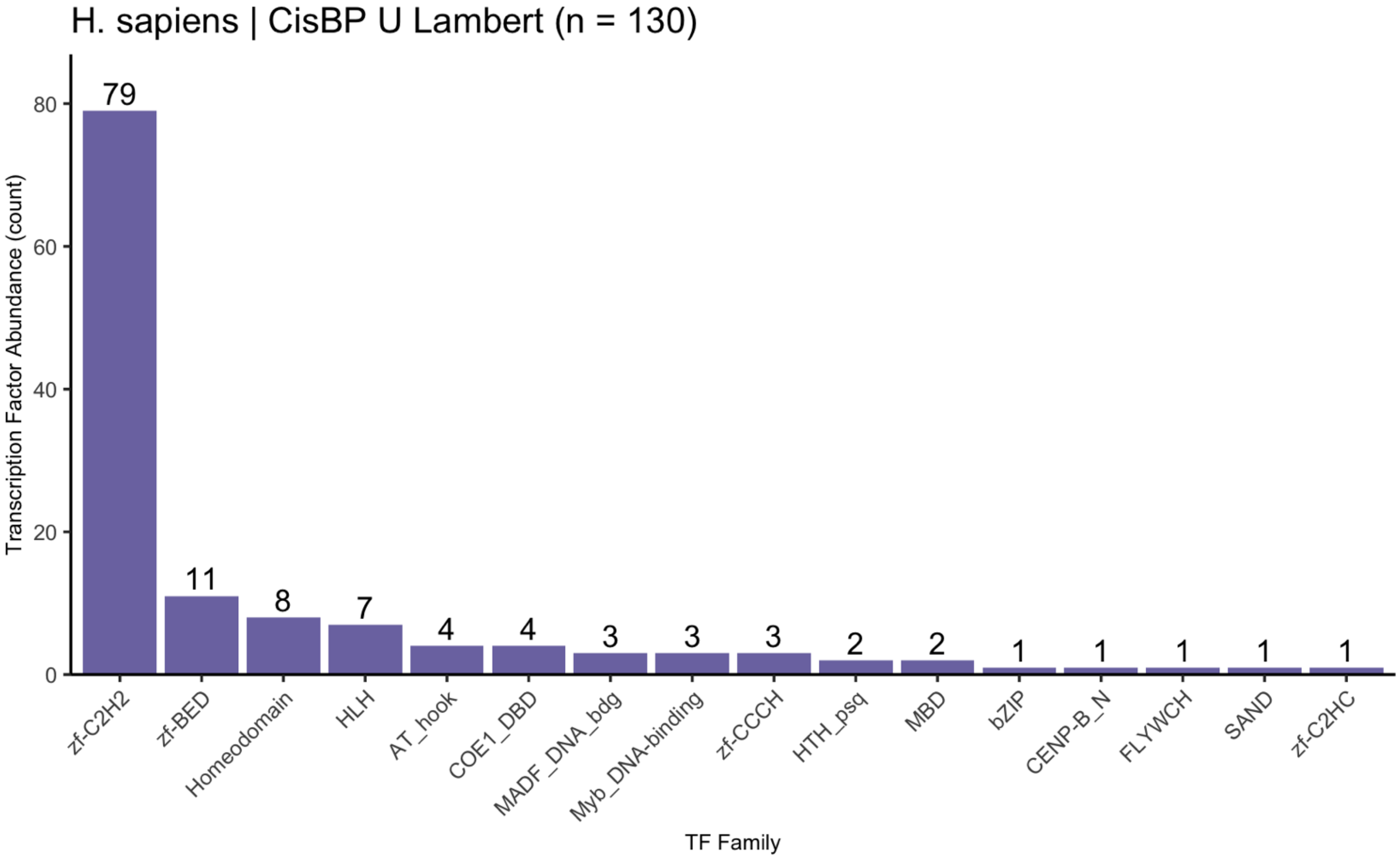


**Supplementary Figure 2:** Transcription factor family distribution plot of the genes not identified by CREPE common to both references.

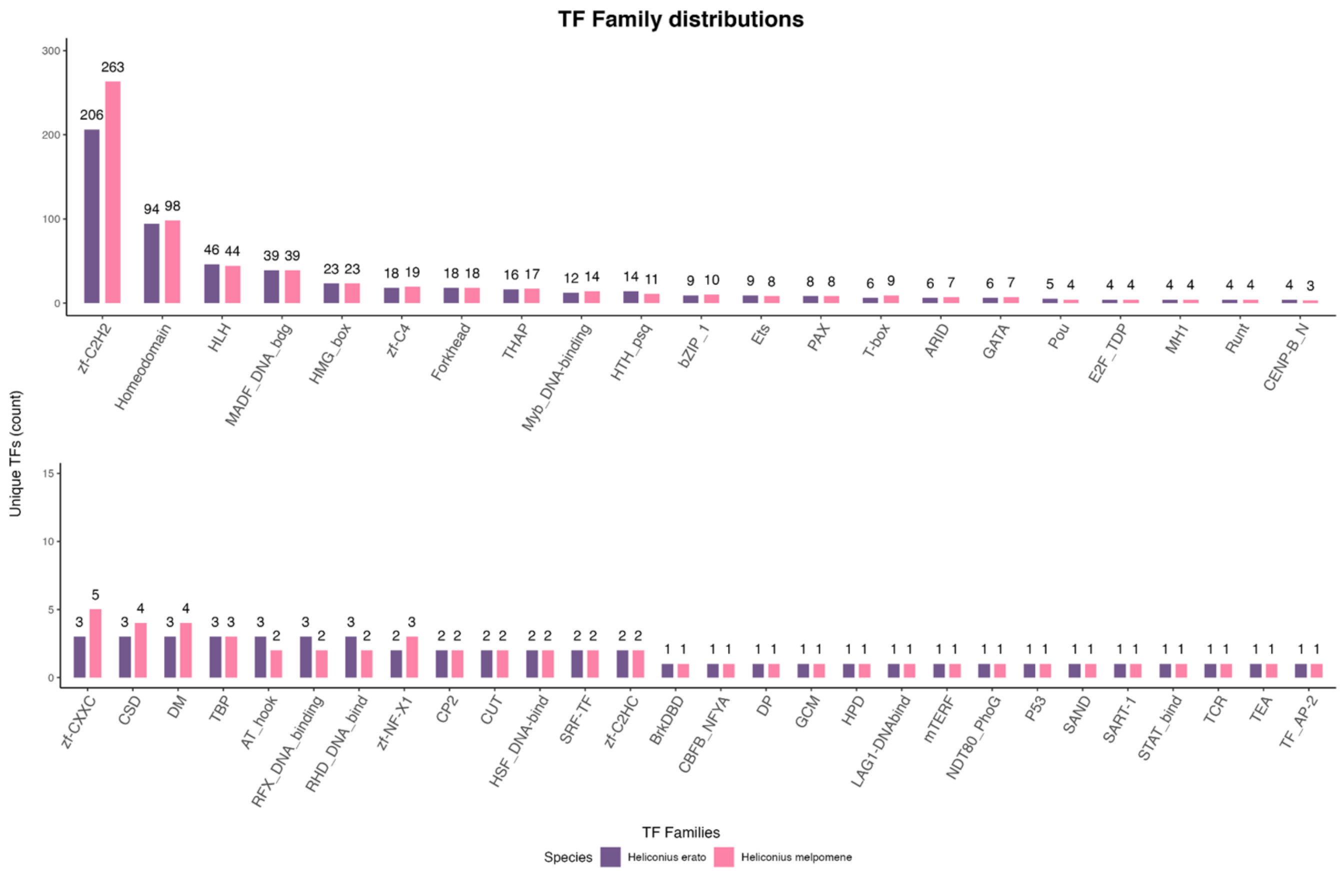


**Supplementary Figure 3:** Transcription factor family distribution in *Heliconius melpomene* and *Heliconius erato*

**Supplementary Table 1:** Summary of the catalogued TFs by family

| index | Pfam_DBDs | CisBP | Lambert | CREPE | Identical |
|---|---|---|---|---|---|
| 1 | zf-C2H2 | 742 | 759 | 681 | N |
| 2 | Homeodomain | 228 | 228 | 228 | Y |
| 3 | HLH | 109 | 108 | 108 | N |
| 4 | bZIP | 54 | 54 | 54 | Y |
| 5 | Forkhead | 49 | 49 | 52 | N |
| 6 | zf-C4 | 46 | 46 | 46 | Y |
| 7 | HMG_box | 29 | 30 | 50 | N |
| 8 | Ets | 28 | 28 | 30 | N |
| 9 | AT_hook | 5 | 25 | 1 | N |
| 10 | T-box | 17 | 17 | 17 | Y |
| 11 | Myb_DNA-binding | 15 | 16 | 25 | N |
| 12 | Pou | 14 | 16 | 16 | N |
| 13 | THAP | 12 | 12 | 13 | N |
| 14 | zf-BED | 12 | 12 | 0 | N |
| 15 | zf-CXXC | 12 | 12 | 13 | N |
| 16 | CENP-B_N | 11 | 11 | 13 | N |
| 17 | E2F_TDP | 11 | 11 | 11 | Y |
| 18 | MBD | 11 | 11 | 9 | N |
| 19 | GATA | 10 | 10 | 14 | N |
| 20 | RHD_DNA_bind | 10 | 10 | 10 | Y |
| 21 | IRF | 9 | 9 | 10 | N |
| 22 | MH1 | 9 | 9 | 12 | N |
| 23 | PAX | 9 | 9 | 9 | Y |
| 24 | RFX_DNA_binding | 9 | 9 | 9 | Y |
| 25 | SAND | 9 | 9 | 9 | Y |
| 26 | HSF_DNA-bind | 8 | 8 | 10 | N |
| 27 | ARID | 7 | 7 | 15 | N |
| 28 | CUT | 7 | 7 | 7 | Y |
| 29 | DM | 7 | 7 | 7 | Y |
| 30 | STAT_bind | 7 | 7 | 7 | Y |
| 31 | CP2 | 6 | 6 | 6 | Y |
| 32 | SRF-TF | 6 | 6 | 6 | Y |

| | | | | | |
|---|---|---|---|---|---|
| 33 | CSD | 5 | 5 | 8 | N |
| 34 | TF_AP-2 | 5 | 5 | 5 | Y |
| 35 | COE1_DBD | 4 | 4 | 0 | N |
| 36 | GTF2I | 4 | 4 | 5 | N |
| 37 | mTERF | 4 | 4 | 4 | Y |
| 38 | TEA | 4 | 4 | 4 | Y |
| 39 | MADF_DNA_bdg | 5 | 3 | 0 | N |
| 40 | P53 | 3 | 3 | 3 | Y |
| 41 | Runt | 3 | 3 | 3 | Y |
| 42 | TBP | 3 | 3 | 3 | Y |
| 43 | zf-CCCH | 3 | 3 | 0 | N |
| 44 | CG-1 | 2 | 2 | 2 | Y |
| 45 | GCM | 2 | 2 | 2 | Y |
| 46 | HPD | 2 | 2 | 2 | Y |
| 47 | HTH_psq | 2 | 2 | 0 | N |
| 48 | LAG1-DNAbind | 2 | 2 | 2 | Y |
| 49 | NDT80_PhoG | 2 | 2 | 2 | Y |
| 50 | zf-NF-X1 | 2 | 2 | 2 | Y |
| 51 | BrkDBD | 1 | 1 | 1 | Y |
| 52 | CBFB_NFYA | 1 | 1 | 1 | Y |
| 53 | FLYWCH | 1 | 1 | 0 | N |
| 54 | TCR | 1 | 1 | 2 | N |
| 55 | zf-C2HC | 7 | 0 | 7 | N |
| 56 | DP | 0 | 0 | 3 | N |
| 57 | SART-1 | 0 | 0 | 1 | N |

**Supplementary Table 2:** Genes in the CREPE-only group

| index | Gene Name | Ensemble Stable ID |
|---|---|---|
| 1 | ARID1A | ENSG00000117713 |
| 2 | ARID1B | ENSG00000049618 |
| 3 | ARID4A | ENSG00000032219 |
| 4 | ARID4B | ENSG00000054267 |
| 5 | CARHSP1 | ENSG00000153048 |
| 6 | CERS2 | ENSG00000143418 |
| 7 | CERS3 | ENSG00000154227 |
| 8 | CERS4 | ENSG00000090661 |
| 9 | CERS5 | ENSG00000139624 |
| 10 | CERS6 | ENSG00000172292 |
| 11 | CPHXL | ENSG00000283755 |
| 12 | CPHXL2 | ENSG00000284484 |
| 13 | CSDC2 | ENSG00000172346 |
| 14 | CSDE1 | ENSG00000009307 |
| 15 | DNAJC1 | ENSG00000136770 |
| 16 | DNAJC2 | ENSG00000105821 |
| 17 | DUXB | ENSG00000282757 |
| 18 | ERFL | ENSG00000268041 |
| 19 | FOXL3 | ENSG00000248767 |
| 20 | FOXO3B | ENSG00000240445 |
| 21 | HMGB1 | ENSG00000189403 |
| 22 | HMGB1P1 | ENSG00000124097 |
| 23 | HMGB2 | ENSG00000164104 |
| 24 | HMGB3 | ENSG00000029993 |
| 25 | HMGB4 | ENSG00000176256 |
| 26 | HMGXB3 | ENSG00000113716 |
| 27 | HMGXB4 | ENSG00000100281 |
| 28 | HSFX3 | ENSG00000283697 |
| 29 | HSFX4 | ENSG00000283463 |
| 30 | ID1 | ENSG00000125968 |
| 31 | ID2 | ENSG00000115738 |
| 32 | ID3 | ENSG00000117318 |

| | | |
|---|---|---|
| 33 | ID4 | ENSG00000172201 |
| 34 | JARID2 | ENSG00000008083 |
| 35 | KDM5A | ENSG00000073614 |
| 36 | KDM5C | ENSG00000126012 |
| 37 | KDM5D | ENSG00000012817 |
| 38 | KLF18 | ENSG00000283039 |
| 39 | KRBOX5 | ENSG00000197302 |
| 40 | MEIOSIN | ENSG00000237452 |
| 41 | MIS18BP1 | ENSG00000129534 |
| 42 | MTA1 | ENSG00000182979 |
| 43 | MTA2 | ENSG00000149480 |
| 44 | MTA3 | ENSG00000057935 |
| 45 | NCOR1 | ENSG00000141027 |
| 46 | NCOR2 | ENSG00000196498 |
| 47 | NFILZ | ENSG00000268480 |
| 48 | NSD2 | ENSG00000109685 |
| 49 | PBRM1 | ENSG00000163939 |
| 50 | PINX1 | ENSG00000258724 |
| 51 | PMS1 | ENSG00000064933 |
| 52 | RCOR1 | ENSG00000089902 |
| 53 | RCOR2 | ENSG00000167771 |
| 54 | RCOR3 | ENSG00000117625 |
| 55 | RERE | ENSG00000142599 |
| 56 | SAMD11 | ENSG00000187634 |
| 57 | SART1 | ENSG00000175467 |
| 58 | SMAD2 | ENSG00000175387 |
| 59 | SMAD6 | ENSG00000137834 |
| 60 | SMAD7 | ENSG00000101665 |
| 61 | SMARCC1 | ENSG00000173473 |
| 62 | SMARCC2 | ENSG00000139613 |
| 63 | SMARCE1 | ENSG00000073584 |
| 64 | SSRP1 | ENSG00000149136 |
| 65 | TADA2A | ENSG00000276234 |
| 66 | TADA2B | ENSG00000173011 |
| 67 | TESMIN | ENSG00000132749 |

| | | |
|---|---|---|
| 68 | TFAM | ENSG00000108064 |
| 69 | TOX | ENSG00000198846 |
| 70 | TOX2 | ENSG00000124191 |
| 71 | TOX3 | ENSG00000103460 |
| 72 | TOX4 | ENSG00000092203 |
| 73 | TPRX2 | ENSG00000259009 |
| 74 | UBTF | ENSG00000108312 |
| 75 | UBTFL1 | ENSG00000255009 |
| 76 | ZFP91-CNTF | ENSG00000255073 |
| 77 | ZKSCAN8P1 | ENSG00000226314 |
| 78 | ZNF286A-TBC1D26 | ENSG00000255104 |
| 79 | ZNF559-ZNF177 | ENSG00000270011 |
| 80 | ZNF722 | ENSG00000241149 |
| 81 | ZNF723 | ENSG00000268696 |
| 82 | ZNF738 | ENSG00000172687 |
| 83 | NA | ENSG00000254553 |
| 84 | NA | ENSG00000289700 |
| 85 | NA | ENSG00000289685 |
| 86 | NA | ENSG00000285708 |
| 87 | NA | ENSG00000283321 |
| 88 | NA | ENSG00000257184 |
| 89 | NA | ENSG00000289346 |
| 90 | NA | ENSG00000284691 |
| 91 | NA | ENSG00000285827 |
| 92 | NA | ENSG00000273049 |
| 93 | NA | ENSG00000258064 |
| 94 | NA | ENSG00000261459 |
| 95 | NA | ENSG00000288000 |
| 96 | NA | ENSG00000283201 |
| 97 | NA | ENSG00000267022 |
| 98 | NA | ENSG00000269693 |
| 99 | NA | ENSG00000196826 |
| 100 | NA | ENSG00000286098 |
| 101 | NA | ENSG00000286132 |
| 102 | NA | ENSG00000142539 |

| | | |
|---|---|---|
| 103 | NA | ENSG00000269825 |

**Supplementary Table 3:** Assemblies used to generate gene trees using CREPE putative TFs and OrthoFinder:

| index | Species | Assembly Name |
|---|---|---|
| 1 | *Anolis carolinensis* | AnoCar2.0v2 |
| 2 | *Balaenoptera musculus* | mBalMus1.v2 |
| 3 | *Bos taurus* | ARS-UCD1.2 |
| 4 | *Caenorhabditis elegans* | WBcel235 |
| 5 | *Ciona savignyi* | CSAV2.0 |
| 6 | *Crocodylus porosus* | CroPor_comp1 |
| 7 | *Danio rerio* | GRCz11 |
| 8 | *Dasypus novemcinctus* | Dasnov3.0 |
| 9 | *Drosophila melanogaster* | BDGP6.32 |
| 10 | *Felis catus* | Felis_catus_9.0 |
| 11 | *Gallus gallus* | GRCg6a |
| 12 | *Homo sapiens* | GRCh38 |
| 13 | *Mus musculus* | GRCm39 |
| 14 | *Myotis lucifugus* | Myoluc2.0 |
| 15 | *Oryzias latipes* | ASM223467v1 |
| 16 | *Pan paniscus* | panpan1.1 |
| 17 | *Pan troglodytes* | Pan_tro_3.0 |
| 18 | *Parus major* | Parus_major1.1 |
| 19 | *Phascolarctos cinereus* | phaCin_unsw_v4.1 |
| 20 | *Pteropus vampyrus* | pteVam1 |
| 21 | *Rattus norvegicus* | Rnor_6.0 |
| 22 | *Saccharomyces cerevisiae* | R64-1-1 |
